# Pharmacological Tools to Modulate Ordered Membrane Domains and Order-Dependent Protein Function

**DOI:** 10.1101/2025.10.03.680351

**Authors:** Katherine M. Stefanski, Hui Huang, Dustin D. Luu, James M. Hutchison, Nilabh Saksena, Alexander J. Fisch, Thomas P. Hasaka, Joshua A. Bauer, Anne K. Kenworthy, Wade D. Van Horn, Charles R. Sanders

**Author notes:** Corresponding authors. (C.S.), (K.S.).

## Abstract

Ordered membrane nanodomains colloquially known as “lipid rafts” have many proposed cellular functions. However, pharmacological tools to modulate protein affinity for rafts and to manipulate raft formation are currently lacking. We screened 24,000 small molecules for compounds that impact the raft affinity of a known raft-preferring protein, peripheral myelin protein 22 (PMP22), in giant plasma membrane vesicles (GPMVs). Hits were counter-screened against another raft protein, MAL, and also tested for their impact on raft stability. We identified three chemically distinct tools for manipulating lipid rafts. Two compounds were seen to both decrease PMP22 raft partitioning and to destabilize ordered domains (VU0607402 and VU0519975) while a third (primaquine diphosphate) increased PMP22 partitioning and stabilized ordered domains. While discovered in a PMP22-focused screen, all three were seen to modulate raft formation in a protein-independent manner by altering lipid-lipid interactions and membrane fluidity. Acute treatment of live cells with the raft destabilizing compound, VU0607402 was seen to modulate TRPM8 channel function, highlighting the utility of this compound in live-cell experiments for dissecting the role that membrane order and fluidity play in cell signaling. These compounds provide novel pharmacological tools for probing lipid raft properties and function in biophysical experiments and in living cells.

## Introduction

It has long been hypothesized that cells utilize transiently-ordered membrane domains, sometimes also called “lipid rafts”, as sorting and signaling platforms (1–6). Specific proteins are thought to be regulated by membrane order, facilitating a variety of biological functions. Lipid rafts have been implicated in immune signaling, neural development, host-pathogen interactions, caveolae biogenesis, cytoskeletal-membrane contacts, and other cellular phenomena, making them a potential therapeutic target (7–12). It is thus desirable to have pharmacological agents to either modulate protein partitioning into rafts or to alter lipid raft formation or to regulate raft-dependent processes. Unfortunately, however, beyond non-specific approaches such as manipulating cholesterol or sphingolipids (13, 14), few methods to pharmacologically target rafts in a controlled manner currently exist.

Under physiological conditions lipid rafts in cells are nanoscale and highly dynamic (15–17). A useful model to study membrane order is giant plasma membrane vesicles (GPMVs). Derived from the plasma membranes of live cells, GPMVs are a well-established tool for studying lipid rafts and for quantitating the raft affinity of membrane proteins (18). GPMVs are easy to prepare and will spontaneously phase separate into micron-scale liquid ordered (L_o_, raft) and liquid disordered (L_d_, non-raft) domains below the miscibility transition temperature. Assorted fluorescent dyes or protein markers are available to label ordered and disordered phases in GPMVs, enabling imaging-based analysis of phases and resident proteins. GPMVs also retain the lipid and protein composition of the cell plasma membrane from which they are derived (19, 20).

GPMVs have also been used to elucidate the characteristics that drive raft affinity of membrane proteins (21–23). The majority of membrane proteins thought to preferentially partition into lipid rafts are single-span transmembrane proteins (21). However, several raft-favoring multispan proteins have also been rigorously identified (22, 24). One example is peripheral myelin protein 22 (PMP22), a tetraspan membrane protein that preferentially partitions into ordered domains in GPMVs (24). PMP22 is thought to play a structural role in myelin compaction and in support of both cholesterol homeostasis and trafficking in Schwann cells of peripheral nerves. Several disease-associated mutant forms of PMP22 exhibit reduced affinity for lipid rafts suggesting the phenomenon is closely related to one of its key physiological functions (24, 25). These considerations motivated us to use PMP22 as a test case to seek compounds that alter protein raft affinity.

In the current study, we used GPMVs as a model system to conduct high throughput screening (HTS) to discover molecules that alter PMP22 partitioning between ordered and disordered domains. Hits from this screen were then tested to see if they also modulate raft formation. Here we describe three hits that increase and decrease phase separation in GPMVs, doing so in a protein-independent manner. The hit compounds were seen to alter membrane fluidity in GPMVs, large unilamellar vesicles (LUVs), and live cells. Finally, we examine the potential for these compounds to modulate biological activities using the human cold-sensing TRPM8 channel as a model system. We show that the raft disrupting compound, VU0607402 reduces TRPM8 signaling. These results identify new tools for investigating raft-dependent phenomena in cells.

## Results

### A high-throughput screen identifies compounds that alter PMP22 raft affinity

To identify compounds that alter the ordered domain partitioning of PMP22, we conducted a high-throughput screen adapted from our recent method (Fig. 1) )(25).

**Figure 1.**
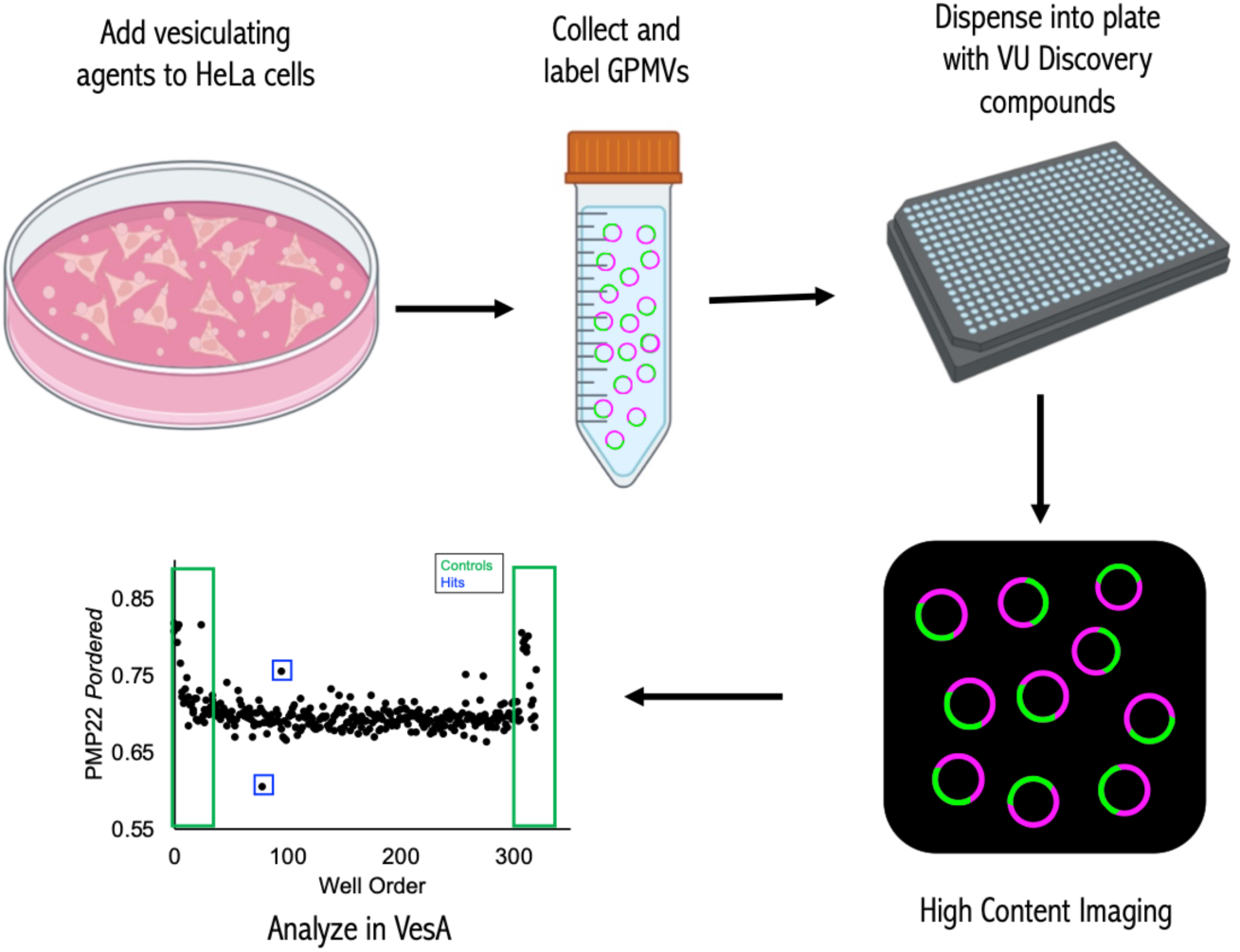
Screening identified modulators of ordered domain affinity. High-throughput screening approach to identify modulators of PMP22 raft affinity. Pipeline used to screen 24,000+ compounds that identified hits described here.

Briefly, the goal was to use GPMVs from cells expressing PMP22 to screen for small molecules that alter the proportion of PMP22 in the ordered phase. GPMVs were made from transfected HeLa cells expressing PMP22. We employed the N41Q (glycosylation deficient) variant of PMP22 since it traffics to the plasma membrane with much higher efficiency than wild type (WT), enabling easier detection, but exhibiting the same ordered domain preference (24, 27). The disordered phase was labeled with the lipophilic stain DiI while PMP22 was immunolabeled with AlexaFluor 647. Next, the labeled GPMVs were used to screen a library of small molecules.

GPMVs were deposited into 96 or 384-well plates containing compounds at a final concentration of 10 µM. A 23,360 compound sub-set of the >100,000 compound Vanderbilt Discovery Collection and a library of FDA approved drugs (1,184) were screened. The Discovery Collection compounds represent a cross-section in terms of the chemical diversity of the master library. Plates were imaged using a high-content confocal imaging system. VesA software was used to analyze images (26). The power of this approach is in the number of vesicles that can be imaged and analyzed in an unbiased manner. A typical well results in images containing thousands of GPMVs which are then analyzed automatically, independent of investigator bias or foreknowledge. Hits from the screen were picked using strictly standardized mean difference (SSMD) values on a plate-by-plate basis (typical cutoff was an SSMD value 290% of the positive control) to assess compound impact on PMP22 *P_ordered_* (28). *P_ordered_* is the fraction of protein in the ordered phase and is a measure of the affinity of a particular protein for rafts. An initial hit rate of just over 1% was observed, resulting in 267 preliminary hits. Hits were then confirmed in triplicate experiments using compounds from the library. Compounds of interest were then reordered from a commercial manufacturer and their effects confirmed. Validated compounds were then tested in GPMVs from cells expressing the MAL protein, another lipid raft-preferring tetraspan protein (22), to test whether their activity is protein-specific. Finally, results were verified in GPMVs from a second cell type, rat basophilic leukemia (RBL) cells, to test for cell type specificity.

The effects of all hits on raft formation were also measured. Here we used the fraction of phase-separated GPMVs (versus non-phase-separated) in our images as a measure of raft formation. For all confirmed compound hits (above), we also performed measurements in GPMVs from untransfected cells. Hits fell into several distinct categories based on these criteria. In this work, we focus on one functionally distinct class of hit compounds.

### Hit compounds alter ordered partitioning of proteins and membrane order

The first criterion for hit picking was a change in PMP22 P_ordered_ and this is the initial basis for the discovery of the compounds (VU0519975, VU0607402 and primaquine diphosphate [PD]) described here. These three compounds are chemically distinct (Table 1). Compounds were then re-ordered and re-tested. Here, two of the compounds, VU0519975 and VU0607402, still showed a decrease in PMP22 *P_ordered_* as in the initial screen, however the effect was not significant while PD maintained a significant effect on increasing PMP22 *P_ordered_* (Fig. 2A). Next, we counter screened the compounds against MAL. A decrease in *P_ordered_* for MAL was induced by VU0519975 and VU0607402 but was not statistically significant (Fig. S1A) while PD induced a significant increase in MAL *P_ordered_* (Fig. S1B).

**Figure 2.**
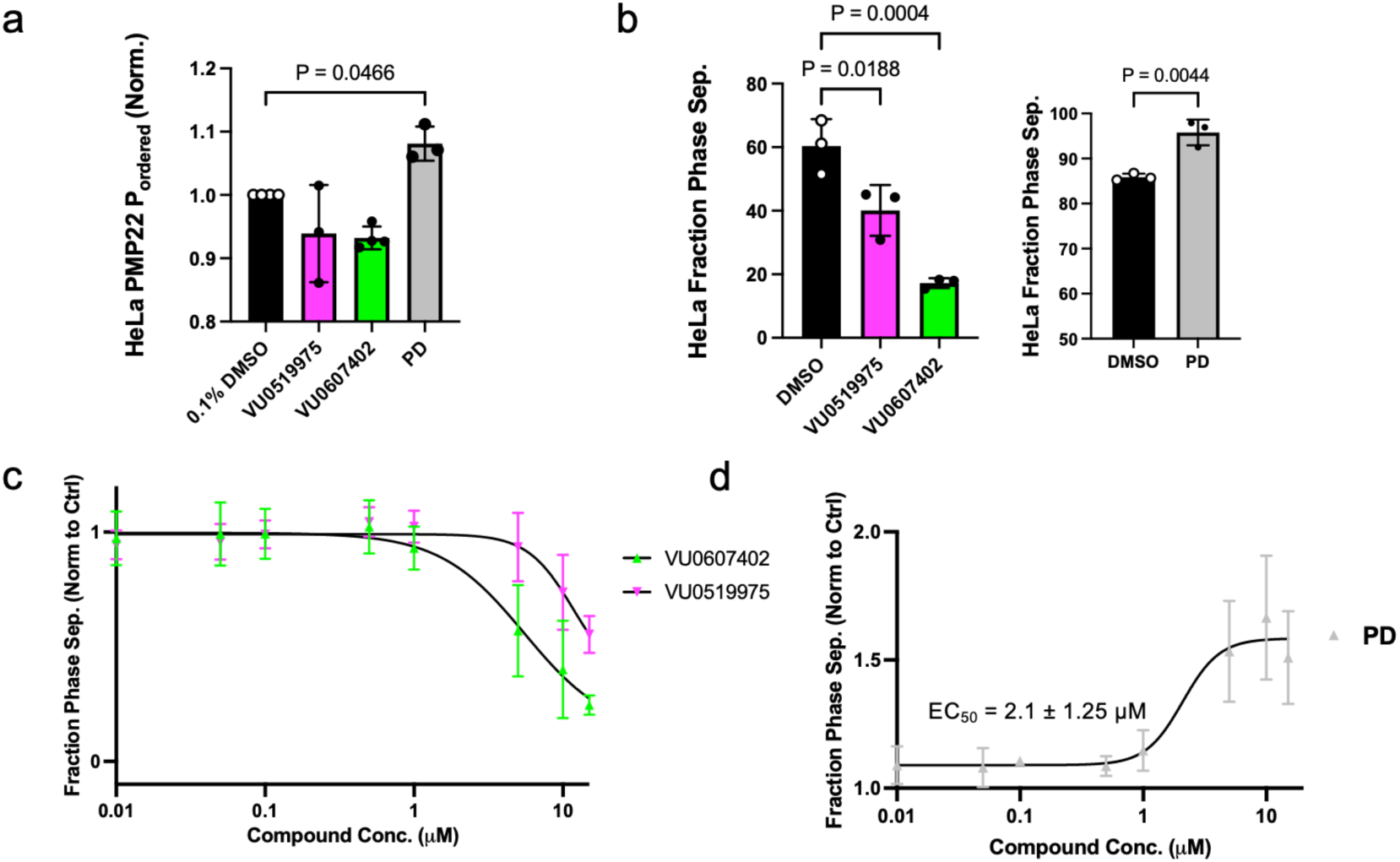
Hit compounds affect ordered partitioning of proteins and ordered domain stability. **a)** Effects of 10 μM VU0519975, VU0607402, and primaquine diphosphate (PD) on ordered partitioning of PMP22 in GPMVs derived from HeLa cells. Bars are means ± SD (*n* = 3), p-values are from Dunnett’s test. **b)** Effects of 10 μM VU0519975, VU0607402, and PD on phase separation with in GPMVs from untransfected HeLa cells. Bars are means ± SD (*n* = 3) **c)** Dose response experiments with VU0519975 and VU0607402 (*n* = 3) **d)** Dose response experiments with PD on phase separation. Points are means ± SD (*n* = 3). Curve and EC_50_ value from sigmoidal fits ± SE.

**Table 1.**
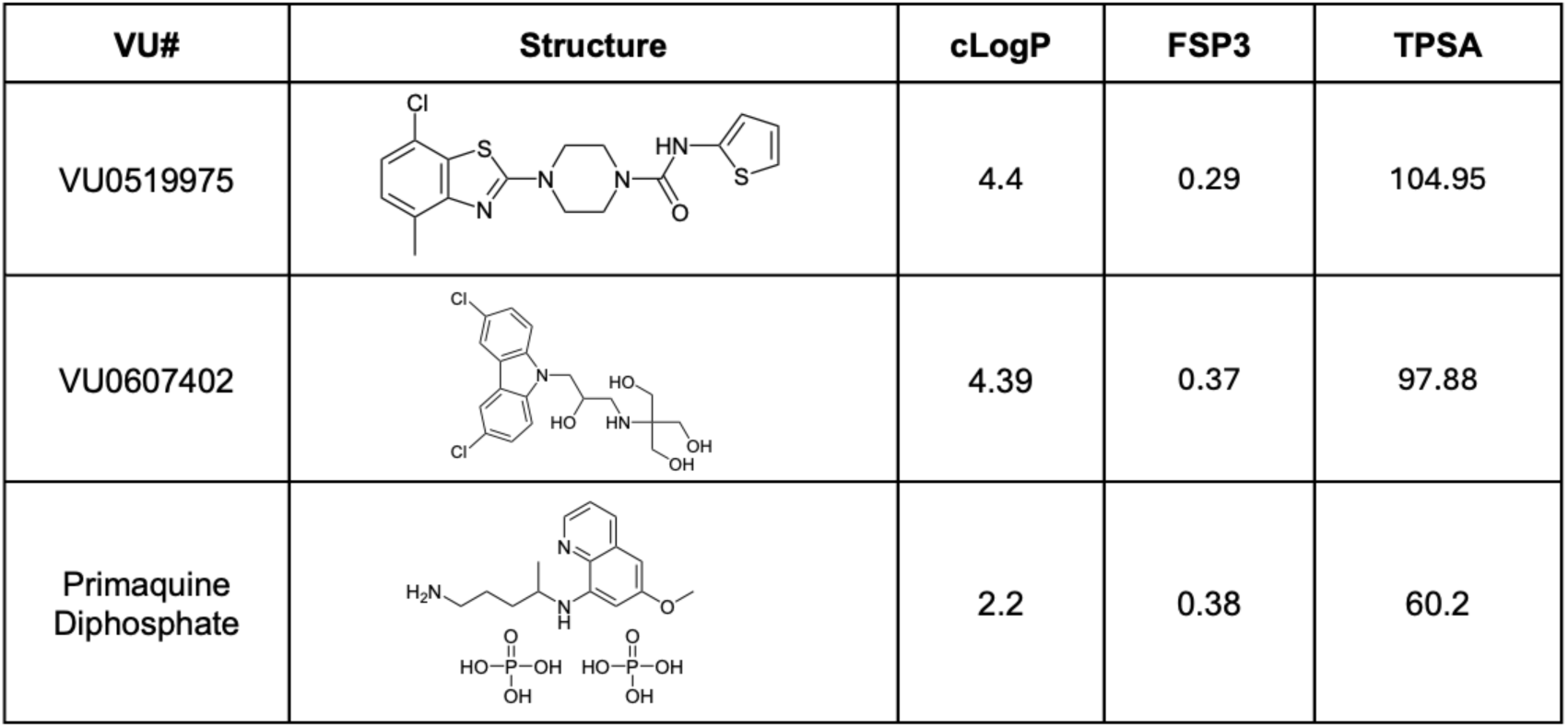
Compound structures and parameters.

| VU# | Structure | cLogP | FSP3 | TPSA |
| --- | --- | --- | --- | --- |
| VU0519975 |  | 4.4 | 0.29 | 104.95 |
| VU0607402 |  | 4.39 | 0.37 | 97.88 |
| Primaquine Diphosphate |  | 2.2 | 0.38 | 60.2 |

When we examined the results of treatment with these compounds on lipid raft stability we found robust effects for all three compounds. Here we used the fraction of phase separated vesicles counted in images of GPMVs as a proxy for raft stability. VU0519975 and VU0607402 significantly decreased the fraction of phase separated GPMVs in images while PD induced a significant increase in the fraction of phase separated GPMVs (Fig. 2B). EC_50_ values for the impact of VU0519975 and VU0607402 on raft stability in GPMVs appear to be in the 5-15 µM range (Fig. 2C) but could not be more accurately determined for since they were insoluble in GPMV buffer at concentrations above 15 µM. EC_50_ for the effect of PD on raft stability in GPMVs was determined to be 2.1 ± 1.25 µM in HeLa derived GPMVs (Fig. 2D).

Notably, these effects were the same in GPMVs made from untransfected cells or from cells expressing PMP22 (Fig S2A). Similarly, there was no effect of MAL expression on the degree to which these compounds destabilize or stabilize lipid rafts (Fig. S1C). We conclude from these data that the compounds exert their effects on membranes independent of either protein examined here and thus likely work through manipulating lipid-lipid interactions.

Finally, we examined the effect of cell type on the effects of these compounds on modulating PMP22 *P_ordered_* and raft stability. To this point, all experiments were conducted in HeLa derived GPMVs that phase separate at temperatures easy to achieve using a variety of instruments (∼25 °C). HeLa cells are also relatively easy to transfect. Another cell line commonly used to form phase-separated GPMVs are rat basophilic leukemia (RBL) cells, which exhibit lower levels of phase separation than HeLa derived GPMVs. We tested the effects of the compounds in GPMVs derived from RBL cells. Here, as expected, we observed that base-line levels of phase separation were lower (Fig S2B, ∼30% phase separated vs 60-80% from HeLa). We found that the compounds VU0519975 and VU0607402 were not soluble in GPMV buffer with RBL cells at 10 µM and so they were tested then at 5 µM. Here we observed that a decrease in raft stability induced by VU0519975 and VU0607402 that was not significant while PD did have a statistically significant effect on increasing raft stability. The more limited effect of VU0519975 and VU0607402 is likely due to the lower concentration used and because the potential effect size was already decreased by the limited phase separation in RBL GPMVs. PD, however, more than doubled the fraction of phase separated RBL GPMVs. Interestingly, VU0519975 and VU0607402 did significantly decrease *P_ordered_* for PMP22 in RBL GPMVs, while PD did not have a significant effect (Fig. S2C).

Neither VU0519975, VU0607402, nor PD affect GPMV size or the relative size of ordered domains in GPMVs (fig. S3 A-D). However, PD induced a small but significant decrease in the size of containing ordered domains in PMP22-expressing cells (figs. S3D-bottom panel and S3E).

### Compounds alter membrane fluidity in vesicles and live cells

To shed light on how these compounds impact raft stability, we next examined how they affect the biophysical properties of membranes, as summarized in Table 2. First, we studied their effects on membrane fluidity. Order and fluidity are closely related physical properties of membranes. Like lipid rafts, membrane fluidity is actively studied for its role in disease and cellular homeostasis (29, 30). Because fluidity experiments can be conducted in model membranes and live cells, this approach also allowed us to connect the effects of the compounds in GPMVs, model membranes, and the plasma membranes of living cells. We used the environmentally sensitive dye Di-4-ANEPPDHQ (Di-4) to report on membrane fluidity (31). Increased membrane fluidity causes a red-shift of the Di-4 emission spectrum, whereas blue-shifted spectra arise from decreased fluidity.

**Table 2.** Summary of biophysical effects of three compounds.

| Compound | Impact on $P_{\text{ordered, PMP22}}$ | Impact on $P_{\text{ordered, MAL}}$ | Impact on raft formation | Impact on raft formation modified by proteins? |
| --- | --- | --- | --- | --- |
| VU0519975 | ↓ (cell type dependent)<sup>a</sup> | no change | ↓ | No |
| VU0607402 | ↓ (cell type dependent)<sup>a</sup> | no change | ↓ | No |
| Primaquine Diphosphate | ↑ (cell type dependent)<sup>b</sup> | ↑ | ↑ | No |

a. A statistically significant decrease is clear in GPMVs RBL cells but not in HeLa cells
b. A statistically significant increase is seen in GPMVs from HeLa cells but not RBL cells

We first treated GPMVs derived from HeLa cells with each compound plus Di-4 and then measured emission spectra. For the two raft destabilizing compounds (VU0519975 and VU0607402), a pronounced redshift was observed in the emission spectra (Fig. 3A). To quantify these changes, generalized polarization (GP) values were determined (Fig. 3B). GP values provide a relative means of comparing spectral shifts. For Di-4 emission, lower values correspond to a more fluid (red-shifted) environment. Compared to DMSO control, the two raft destabilizing compounds significantly decreased measured GP values indicating that they significantly increase membrane fluidity. However, PD did not significantly affect fluidity in these experiments.

**Figure 3.**
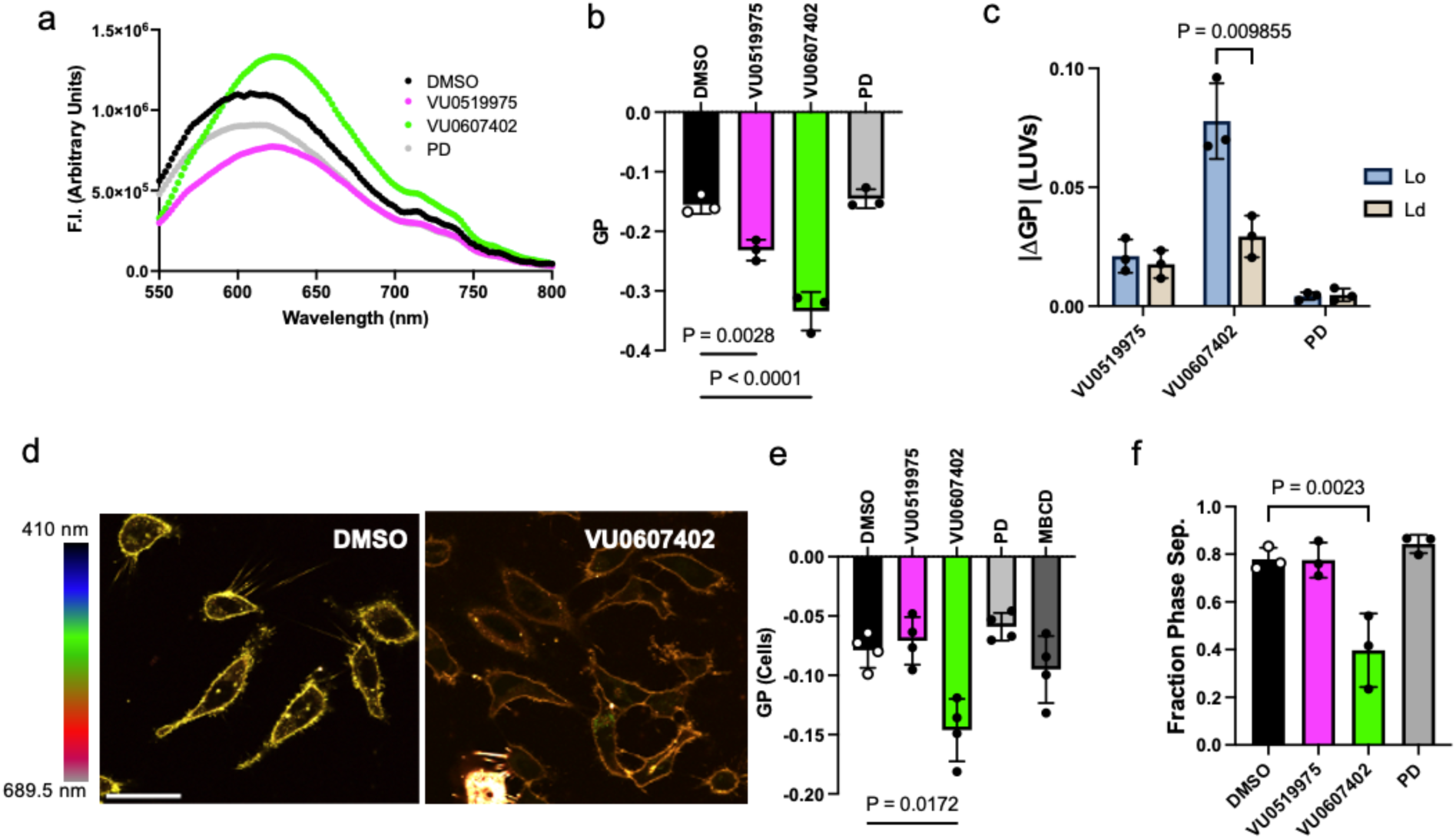
Compounds alter membrane fluidity in GPMVs, live cells and LUVs. **a)** Representative Di-4 emission spectra of GPMVs treated with 10 μM hit compounds. **b)** Generalized polarization values calculated from Di-4 emission spectra shown in a. Bars are means ± SD (*n* = 3). P-values are from ANOVA followed by Dunnett’s multiple comparisons tests. **c)** Absolute value of change in GP from DMSO control in Lo and Ld LUVs. P-values are from unpaired student’s t-tests, *n* = 3. **d)** Representative spectral images of live HeLa cells stained with Di-4 and treated with DMSO or 10 μM VU0607402. Scale bar = 50 μm. **e)** Generalized polarization values calculated from Di-4 emission intensities calculated from individual cells as shown in c. Bars are means ± SD, 10-15 cells (technical replicates) per treatment were measured for each of 4 biological replicates. P-values are from ANOVA followed by Dunnett’s multiple comparisons tests. **e)** Effects compounds on phase separation in GPMVs from untransfected HeLa cells pre-treated with 10 μM compound prior to vesiculation. Bars are means ± SD (*n* = 3) P-values are from ANOVA followed by Dunnett’s multiple comparisons tests.

GPMVs consist of both protein and lipid components. To gain mechanistic insights into the effects of the compounds on lipid bilayers, we performed Di-4 fluidity experiments in large unilamellar vesicles (LUVs). We generated LUVs consisting of mixtures of POPC, sphingomyelin (SM), and cholesterol at concentrations known to form either uniform liquid ordered (L_o_, 25% POPC, 35% SM, 40% cholesterol) or liquid disordered (L_d_, 70% POPC, 25% SM, 5% cholesterol) vesicles (32). We then examined the absolute difference in GP values with and without compound in the L_o_ versus L_d_ LUVs (Fig. 3C). The compounds altered GP in both ordered and disordered LUVs. As in GPMVs, VU0519975 and VU0607402 treatment changed fluidity, while PD had little effect. This indicates VU0519975 and VU0607402 enhance membrane fluidity in lipid-only bilayers and supports the notion that the compounds work through promoting or disrupting lipid-lipid interactions. Interestingly, the effects of VU0607402 were significantly larger in the ordered LUVs compared to the disordered LUVs suggesting that this compound interacts with membranes in a different manner than VU0519975 and PD, and that its activity is sensitive to membrane composition.

Next, we assessed the effects of the compounds on membrane fluidity in living cells. HeLa cells were treated with compounds ∼15 min before a brief incubation with Di-4 to limit internalization of the dye and ensure primarily plasma membrane labeling. Cells were then imaged via laser scanning confocal microscopy using a spectral detector, allowing images from multiple emission wavelengths to be simultaneously acquired (Fig. 3D and fig. S4A). GP values were then calculated from the images. We found that the changes in GP were very similar to those from GPMVs (Fig. 3E). VU0607402 produced statistically significant increases in membrane fluidity. While not statistically significant, the effects of VU0519975 and PD on Di-4 emission in live cells show similar changes relative to each other as was observed in the GPMV Di-4 experiments (Fig. 3, compare B and E).

To confirm that the changes in membrane fluidity observed in live cells were related to the effects of the compounds on plasma membrane rafts, we performed an experiment in which we isolated GPMVs from cells that had been pre-treated with each compound. We found that GPMVs derived from cells treated with VU0607402 exhibited decreased phase separation compared to controls (Fig. 3F). This mirrors the effect of VU0607402 on phase separation observed in the original screen in which the GPMVs were treated directly with the compounds (Fig. 1B). In contrast, no difference in phase separation was observed in GPMVs isolated from cells pretreated with VU0519975 or PD compared to control GPMVs (Fig. 3F). These findings suggest (i) that the increase in membrane fluidity and raft disruption in response to VU0607402 treatment in living cells are coupled and (ii) PD and VU0519975 are either rapidly metabolized by cells or are unable to overcome other raft stabilizing forces in intact plasma membranes.

To compare the effects of our compounds with that of cholesterol depletion, we added methyl-β-cyclodextrin (MBCD) to HeLa cells to deplete membranes of cholesterol. In comparison to the effects of VU0607402, the effect of MBCD treatment on membrane fluidity was modest. Qualitatively, the MBCD-treated cells also appeared rounded (fig. S4A, bottom row) compared to the compound-treated cells, which retained the well-spread appearance of untreated cells. Trypan blue cell viability experiments confirmed that MBCD shows toxicity in cells while none of the 3 compounds reported here were cytotoxic at the concentrations and timescales used in the live cell experiments (fig. S4B). This suggested these compounds are better tolerated by live cells than MBCD and could serve as improved tools for manipulating membrane order in cells to study the functional relevance of lipid rafts in different biological processes. There are no PMP22 or MAL functional assays. We therefore next investigated their impact on functions of other membrane proteins previously proposed to be raft dependent in live cells.

### VU0607402 alters TRPM8 channel activity but not autophosphorylation of EGFR

To test if these compounds impact lipid raft-dependent processes in cells, we examined their effects on the cold and menthol sensing ion channel human TRPM8 (33, 34) as a test case. Many ion channels, including TRPM8, are thought to associate with lipid rafts and have signaling properties that are sensitive to membrane order (35–37). We chose two compounds for these studies: VU0607402 and PD. VU0607402 was selected based on its ability to exert significant effects on cell membrane fluidity, while PD was of interest since it has the opposite effect on raft stability in model membranes compared to VU0607402.

We conducted automated patch clamp (APC) electrophysiology experiments (Fig. 4A, fig. S5A), a technique that allows for prolonged patching of cells, to determine if compound treatment altered TRPM8 function. TRPM8-expressing cells were equilibrated to 30 °C (38) and treated with the canonical TRPM8 agonist menthol before and after 15 minutes of continuous perfusion with the lipid raft-modulating compounds. The pre- and post-menthol-stimulated currents were compared to evaluate the effects of the raft-modulating compounds on TRPM8 function. MBCD was included for comparison since previously published studies used MBCD to deplete cholesterol and disrupt lipid rafts and examine TRPM8 signaling (38).

**Figure 4.**
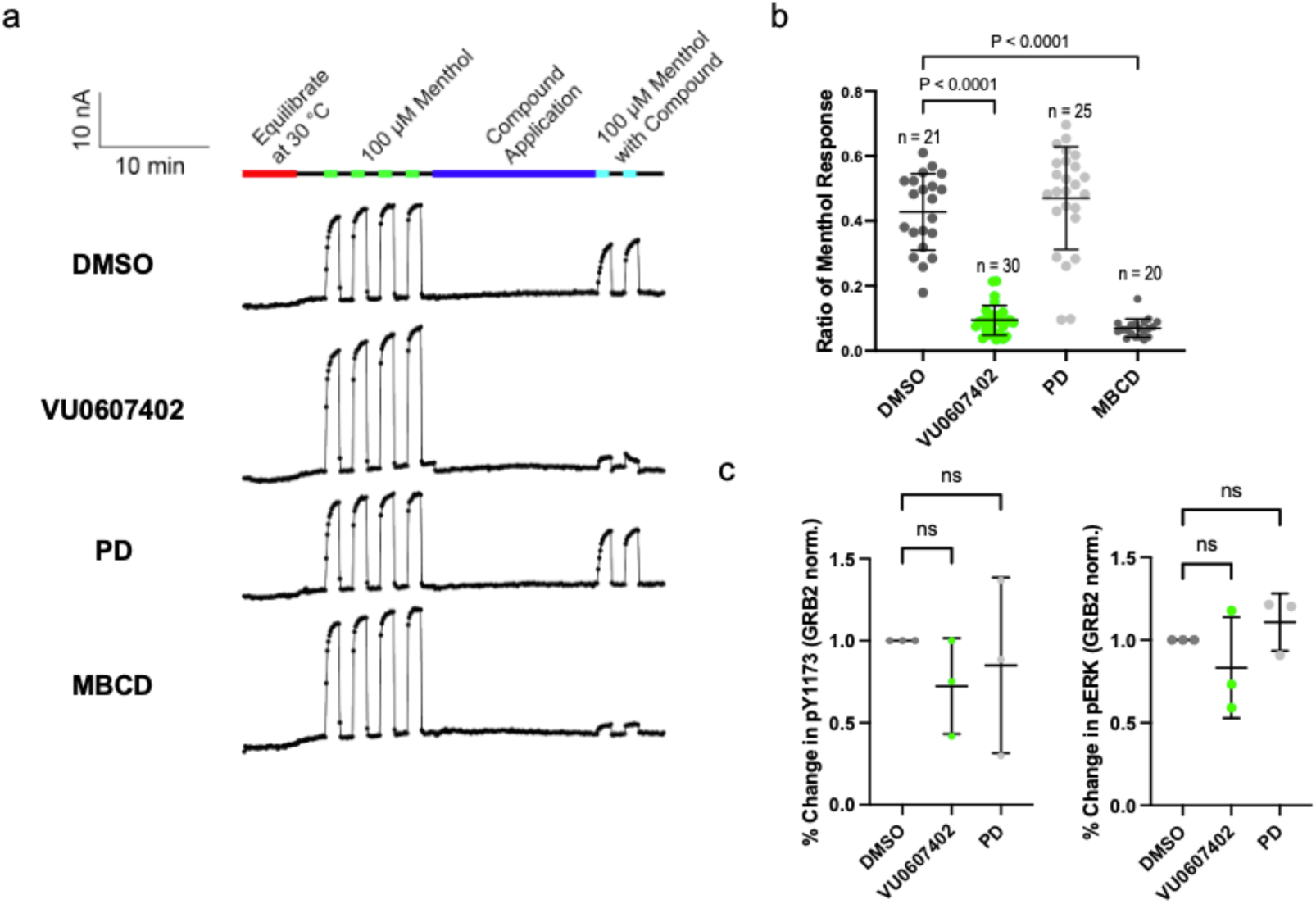
VU0607402 decreases the activity of TRPM8 but not EGFR. **a)** Average current traces of TRPM8 menthol response before and after exposure to the compounds in HEK293T cells expressing human TRPM8 from automated patch clamp experiments. Each dot is when the pulse program is applied. 100 µM menthol was applied for 75 seconds 4 times before perfusing 0.03% DMSO (control) or 10 µM compounds for 15 min. 100 µM menthol was then applied with corresponding compounds twice for 75 seconds. One n is the sum of 20 cells from an amplifier on the plate. **b)** Average ratio of menthol response from each compound in TRPM8 HEK293T cells. The ratio calculated from data in panel a, where the last two menthol responses before compound application were averaged and compared to the two menthol responses after compound application. P-values were determined by ANOVA followed by Dunnett’s tests. Bars are means ± SD. **c)** Quantification of of Western blot analysis of phospho-EGFR (left) and ERK (right) from HeLa cells treated with compounds for 15 min then stimulated with EGF for 1 min. n =3, bars are means ± SD. All comparisons are not significant.

We found that perfusion with VU0607402 and MBCD reduced the TRPM8 current responses to menthol stimulation (Fig. 4B, fig. S5A). These results support the notion that TRPM8 function is dependent on lipid rafts and/or changes in membrane fluidity. PD did not significantly alter TRPM8 activity. These results resemble those seen in the live-cell fluidity experiments and provide important cross-validation.

We considered the possibility that the raft modulators may impact channel function by changing their abundance at the plasma membrane. However, treatment with VU0607402 or PD did not alter cell surface levels of TRPM8 as quantified by flow cytometry (fig. S5B). This suggest VU0607402 is not altering TRPM8 activity by removing the protein from the cell surface. Interestingly, MBCD increased the level of surface TRPM8, likely reflecting its known activity as an inhibitor of endocytosis (39). Together with the membrane fluidity and cell viability results, these data show that VU0607402 has more robust effects on plasma membrane fluidity than MBCD and produces a similar decrease in TRPM8 signaling without changing the levels of TRPM8 at the plasma membrane or inducing cytotoxicity.

As a second test case we examined the impact of raft modulators on the activation of the epidermal growth factor receptor (EGFR) by epidermal growth factor (EGF). Previous studies have shown changes in EGFR activation following disruption of lipid rafts by MBCD treatment (40–42). We treated HeLa cells with VU0607402 or PD for 15 minutes (as in the TRPM8 experiments) prior to a brief (1 min) treatment with EGF. Cells were then lysed and levels of EGFR phospho-tyrosine 1173 and phospho-ERK (phosphorylated downstream of EGFR phosphorylation) were quantified via Western blot analysis. We did not see significant effects on either EGFR or ERK phosphorylation (Fig. 4C, fig. S6), indicating that these signaling processes are not sensitive to short-term changes in membrane fluidity or raft modulation. These data also indicate that not all plasma membrane signaling events are sensitive to the changes in membrane order and fluidity stability induced by these compounds.

## Discussion

Studies of function of lipid rafts have been stymied by the lack of pharmacological tools to systematically manipulate their stability and composition. Here we present 3 new small molecules, VU0519975, VU0607402, and PD, that modulate the partitioning of certain membrane proteins and raft formation at micromolar concentrations. These two activities appear to not be coupled, as the impact of these compounds on raft formation was found to be protein-independent. All three molecules were non-toxic to cells. We identified two raft destabilizers and one raft stabilizer, PD. The compounds exerted effects in both synthetic model membranes and in isolated plasma membrane vesicles. VU0607402 consistently exerted effects in live-cell plasma membranes. VU0607402 altered the signaling of a raft-sensitive ion channel, TRPM8. It also induced a more robust increase in cell membrane fluidity than MBCD. These compounds, especially VU0607402, should be useful pharmacological tools for manipulating lipid raft formation and raft-partitioning of membrane proteins in biophysical and cell biological studies.

The structures of these compounds provide clues into how they likely interact with membranes. All are all moderately hydrophobic, with cLogP values in the 2-4.5 range, below the threshold for near-100% partitioning into the interior of membranes or fat. The molecular architecture of these molecules is not lipid-like. Each compound has aromatic moieties with attached polar groups. Aromatic groups containing a nitrogen (pyridine-like) or with directly attached hydrogen bonding moieties preferentially partition into membranes in the headgroup/interface region, not deep in the membrane interior (43). This observation suggests that all three molecules interact with membranes near the water-bilayer interface, which allows their apolar (mainly aromatic moieties) to extend into the membrane surface below the polar headgroups of the lipids, while their more polar moieties are located in the hydrated headgroup region of the membrane. With this simplistic but reasonable model for how these molecules interact with membranes, we can propose mechanisms for how both types of molecules modulate raft stability.

### Proposed mechanism of action for modulation of raft formation

We propose that PD and VU0519975 influence membrane organization primarily through interactions with lipid headgroups to differentially alter lipid-lipid interactions in the ordered phase versus the disordered phase (Fig. 5). Together with favorable hydrocarbon packing, backbone/headgroup hydrogen bonding between sphingolipids and cholesterol are thought to play a role in stabilizing ordered domains (1, 44, 45). PD (with the lowest cLogP of the compounds described here) has been previously described to interact primarily with phospholipid headgroups (46). We think this explains its protein-independence, limited activity in membrane fluidity experiments, and apparent insensitivity to membrane composition (high cholesterol and SM vs low cholesterol and SM) observed in the experiments in LUV experiments. We propose PD acts by enhancing lipid-lipid interactions and hence, order, favoring the ordered phase.

**Figure 5.**
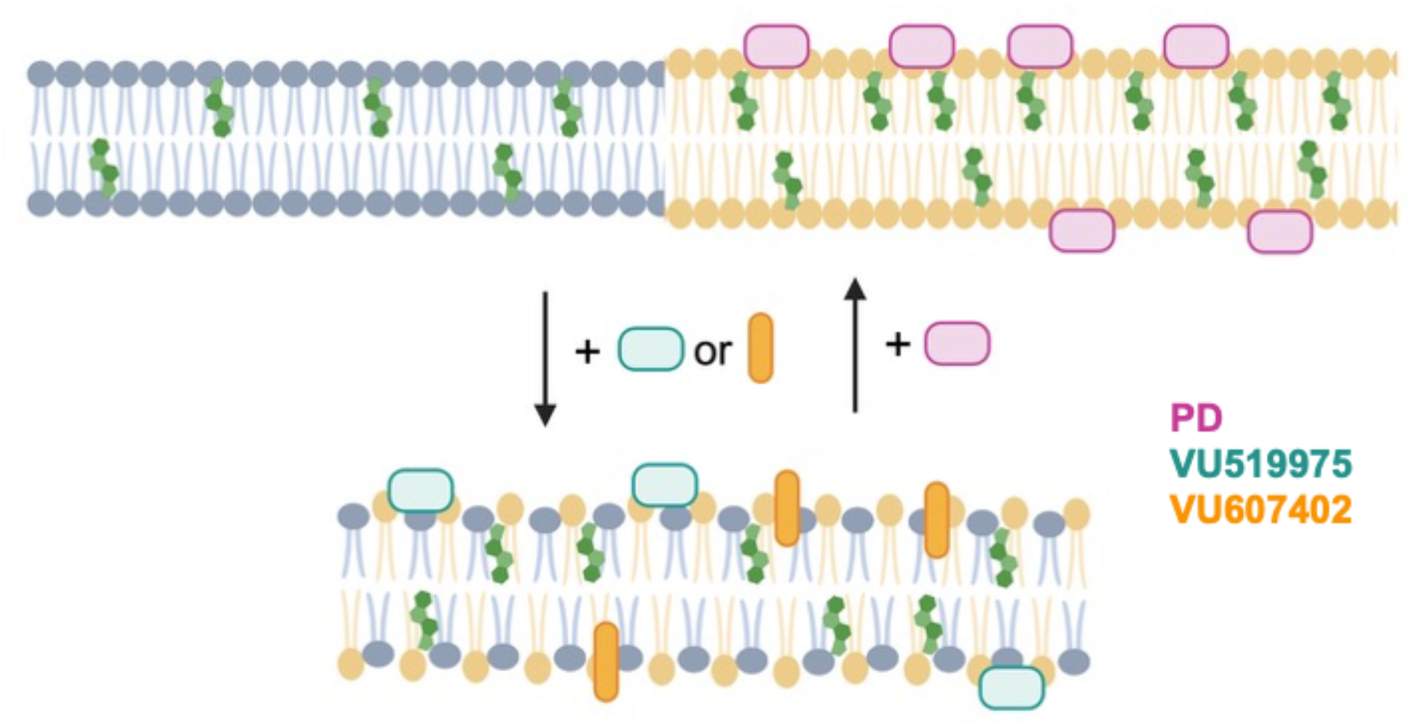
Proposed mechanisms of membrane order modulating compounds. Compounds promote or destabilize phase separation. PD and VU519975 likely interact with lipid headgroups to promote or destabilize lipid-lipid interactions. VU607402 may intercalate into the membrane to promote fluidity via acyl chain interactions.

VU0519975 showed a similar insensitivity to membrane composition as PD in LUV fluidity experiments and may therefore interact with the membrane in a similar manner. However, it had an opposite effect from PD, promoting the disordered phase. We propose that VU0519975 disrupts favorable headgroup/backbone-located lipid-lipid interactions, thereby disfavoring the ordered phase.

VU0607402 also promotes the disordered phase. We propose that VU0607402, unlike the other two compounds, interacts with the membrane via some degree of burial where it promotes disorder and increases fluidity preferentially in ordered membranes (Fig. 5). VU0607402, also known as DCAP, is a broad-spectrum antibiotic that targets bacterial cytoplasmic membranes, perturbing the membrane potential, which leads to mislocalization of proteins important for fission (47). DCAP has also been shown to inhibit autophagy in mammalian cells by blocking autophagolysosome maturation (48). Membrane composition and order have been implicated in autophagosome-lysosome fusion (49). Future studies will be needed to determine if the observed effects of DCAP on bacterial membrane potential or on mammalian autophagy are caused by disruption of membrane order.

### Impact of Compounds on Raft versus Disordered Phase Partitioning of PMP22

While the focus of this work ended up being on the impact of VU0519975, VU0607402, and PD on raft formation, it should not be forgotten that these compounds altered the phase partitioning of PMP22 in GPMVs while P_ordered_ for another raft-preferring membrane protein, MAL, remained statistically unperturbed (see Table 2 for summary). While these are preliminary results and will require further exploration and testing, it suggests that it may be possible to discover protein-specific modulators of raft affinity. This would be a very welcome development for studies of how the function of specific proteins are altered by raft association and potentially even for therapeutic applications.

### Preliminary applications of compounds in membrane biology

Previously used methods for altering lipid rafts in cells include removal or delivery of cholesterol via MBCD and use of alcohols of varying chain lengths (50–52). These approaches have serious limitations. MBCD efficiently removes cholesterol, but this is ultimately lethal to cells. MBCD is also not specific and can remove phospholipids as well as cholesterol (51). Hexadecanol and octanol can be used to decrease or increase membrane fluidity, respectively (50), but they have low miscibility in aqueous buffers and media, making them challenging to work with in cell-based assays. We also recently described bioactive compounds that either promote or reduce raft formation, but the previously reported compounds are all known to have effects on various cellular functions in addition to their membrane effects (26). For example, one is a protease inhibitor. VU0607402 presents as a new and improved tool for lipid raft manipulation in live cells.

Potential applications VU607402 are exemplified in our studies with TRPM8 and EGFR. Prior studies implicated lipid raft localization as having regulatory effects on the activities of both TRPM8 and EGFR (40, 41, 53). Our results showed that micromolar levels of VU0607402, which decreases raft formation in GPMVs, robustly decreased TRPM8 signaling. We note that previous studies reported an *increase* in rat TRPM8 channel activity after treatment with MBCD (38, 54). The difference is likely the result of the speciation difference between the human TRPM8 used in this study compared to the rat TRPM8 used in the previous work (55, 56). Speciation differences have also been seen in various other TRP channels, including TRPA1, TRPV3, and TRPV1 (57, 58). In our experiments, we also observed that MBCD altered the levels of TRPM8 at the cell surface, which must be considered as a confounding variable when interpreting experiments using MBCD.

Reduced raft formation induced by treatment with the same compound, VU607402, that affected TRPM8 had little to no effect on EGFR phosphorylation, in contrast to the reported impact of lipid raft disruption by MCBD treatment (40, 41, 59). This suggests that EGFR activity may be more sensitive to the cholesterol concentration in the membrane than it is to lipid rafts. Examples of membrane proteins that sense and are regulated by specific lipids are replete in the literature. Some G protein-coupled receptors are known to be allosterically regulated by direct binding of cholesterol (60). Another example is the yeast transcriptional regulator Mga2, which demonstrates that some proteins sense acyl chain composition rather than overall fluidity and respond with a conformational change that regulates their activity (61).

### Conclusions

The discovery and characterization of the molecules in this work presents new pharmacological tools for interrogating lipid rafts and raft proteins in cells. Moreover, our initial results using these compounds shed light on the fact that order-stabilizing lipid-lipid interactions can be manipulated independent of proteins.

Further studies will be needed to fully reveal how the compounds described here exert their effects. It seems unlikely they would change cellular lipid composition (as MBCD does) on the short time scales (∼15 min) they were tested in this work, especially in light of their structures. However, future lipidomic analyses could test this possibility. We speculate that longer-term incubation of live cells with the compounds could result in membrane remodeling as cells respond to changes in raft stability and membrane fluidity. Cells have adapted mechanisms to sense and respond to these changes resulting from natural temperature changes, driven by the imperative to keep their membranes near the miscibility critical point (62–65). Further experiments with these compounds can also broaden our understanding of these mechanisms.

## Materials and Methods

### Cell culture

HeLa and RBL-2H3 cells and were acquired from the American Tissue Culture Collection (ATCC, Manassas Va, cat #CCL-2 and CRL-2256). Cells were grown at 37 °C in 5% CO_2_ in a humidified incubator. HeLa cells were cultured in low glucose DMEM (Gibco # 11885084) supplemented with 10% fetal bovine serum (FBS, Gibco, #26140-079) and 100 U/mL penicillin and 100 µg/mL streptomycin (P/S, Gibco, #15140-122). RBL-2H# cells were cultured in MEM (Gibco 11095808) with 10% FBS and 100 U/mL penicillin and 100 µg/mL streptomycin.

### GPMV formation and imaging

HeLa cells were plated at ∼1.4 × 10^6^ cells total in a 150 mm plate. For PMP22 experiments: 24 hrs later cells were transfected with 15 µg of pSF PMP22 N41Q-myc using Fugene 6 (Promega cat # E2691) following the manufacturers protocol. Cells were grown for an additional 48 hrs post transfection. In counter screening experiments, cells were transfected with a MAL-GFP construct (gift from the Levental lab (22)) or transferrin receptor (TfR)-mCherry. Where cells were pre-treated with compound prior to vesiculation, untransfected HeLa cells were incubated with compounds in complete media at 37 °C for 30 minutes. The compounds were then rinsed away prior to GPMV formation. GPMVs were generated using a standard protocol(18), cells were first rinsed twice with 10 ml of inactive GPMV buffer (150 mM NaCl, 10 mM HEPES, 2 mM CaCl2, pH7.4). To label the disordered phase, cells were stained with DiIC12 (3) (1,1’-Didodecyl-3,3,3’,3’-Tetramethylindocarbocyanine Perchlorate) (DiI, Invitrogen cat # D383) for 10 minutes. Cells were then rinsed twice again with inactive GPMV buffer then subsequently incubated in 8.5-10 ml of active GPMV buffer (GPMV buffer + 2 mM DTT, 25 mM formaldehyde) at 37°C for 1.5 hrs. After incubating, GPMVs in solution were collected from the dish and allowed to settle at room temperature for 1 hr. 8-9 mL of GPMVs were then collected by pipetting from neither the top nor the bottom of the tube to leave behind both floating and settled debris. A mouse anti-myc antibody (Cell Signaling, 9B11) was then added at a ratio of 1:1500 and incubated for 1 hr. This was followed by incubation with an anti-mouse antibody conjugated to AlexaFluor647 (Cell Signaling, cat # 4410) at 1:15000. GPMVs were added to multi-well plates with compounds from DMSO stocks and allowed to incubate for 1.5 hrs. Plates were then imaged on an ImageXpress Micro Confocal (IXMC) High Content Screening System (Molecular Devices, San Jose CA) with a Nikon 40X 0.95 NA Plan Apo Lambda objective and an Andor Zyla 4.2MP 83% QE sCMOS camera, and an 89-North LDI 5 channel laser light source.

### High-throughput screen and GPMV image analysis

A pilot screen was conducted with an FDA approved drug library (1,184 compounds) by testing the library plated in triplicate. Primaquine diphosphate (PD) was found to have significant effects on promoting PMP22 ordered partitioning and increasing raft stability. It was carried through the larger screen as a positive control. Compounds were obtained from the Vanderbilt Discovery collection at the VU HTS core. This library contains over 100,000 compounds with the first 20,000 representing the greatest structural diversity. Compounds were dispensed via a Labcyte Echo 555 into 384-well plates such that final the final screening concentration for all compounds was 10 µM. GPMVs were made and labeled as described above and added to the plates containing compounds. The first and last columns of the plates were filled with positive and negative controls. Plates were incubated for at least 1.5 hours at room temperature prior to imaging. 16 images per well were collected using an IXMC as described above.

Images were then analyzed using VesA. Strictly standardized mean differences (SSMDs) were used to calculate effect sizes for the PMP22 ordered phase partition coefficient (*P_ordered_*), for every well (28, 66). The SSMDs for positive (PD) and negative (DMSO) controls were used as a benchmark to select hits from each plate. A typical cutoff for selecting hits was a SSMD value of >90% of the positive control SSMD for PMP22 ordered partitioning. These criteria resulted in a 1-3% hit rate. SSMD values for PMP22 ordered partitioning were used to pick hits. For each hit we also noted the fraction of phase-separated vesicles (also calculated by VesA). A minimum of 100 GPMVs expressing PMP22 per well was required to be included as a hit. Hits were then screened by qualitatively assessing the images from screening. Hits that showed significant visible compound precipitation or extreme changes in GPMV size/shape were discarded. After screening 23,360 compounds 267 hits were identified following the above criteria, reflecting a 1.06% hit rate. These hits were then tested in triplicate against library compound. From hits that were validated at this phase, those with the largest effects were selected and reordered from a commercial vendor (20 compounds, vendor list below).

Experiments using reordered compounds were conducted in 96-well plates with duplicate or triplicate wells (technical replicates) in each plate (biological replicate). For experiments with PMP22 or MAL, GPMVs containing the overexpressed construct were analyzed. For follow-up and dose response experiments, a minimum of 10 phase-separated GPMVs per biological replicate were required for ordered partitioning measurements (determination of *P_ordered_*). For experiments without PMP22 or MAL, (untransfected cells) all GPMVs were analyzed.

### Compound repurchasing

Compounds from the Vanderbilt Discovery Collection were reordered from Life Chemicals (VU0519975, Cat No. F5773-0110) (VU0607402, Cat. No. F3255-0148)

(Niagara on the Lake, Ontario, Canada). PD was acquired from Selleckchem (Cat. No. S4237) (Houston, TX, USA).

### Dose-response experiments

GPMVs from cell expressing PMP22 or MAL or from untransfected cells were prepared and labeled as described above. Doses of hit compounds or DMSO were made by serial dilution and deposited into wells of a 96-well plate for a final concentration from 0.01 to 15 µM. The measured fraction of phase-separated GPMVs and *P_ordered_* were normalized to DMSO controls and averaged. Curves were fit in GraphPad Prism10 and half maximal effective concentrations (EC_50_) were determined using a non-linear sigmoidal model.

### Proteinase K treatment

Protease treatment of GPMVs was conducted as previously described(67). GPMVs were made from untransfected HeLa as described above and labeled with DiD and NBD-PE. GPMVs were then separated into 2 tubes. One tube was treated with 20 µg/mL of proteinase K (Macherey-Nagel cat #740506). Proteinase K and untreated GPMVs were incubated for 45 min at 37° C. 2 mM PMSF was added to quench the proteinase K treatment. GPMVs were then added to wells in a 96-well plate containing compounds for a final concentration of 10 µM. GPMVs were imaged and analyzed as described above.

### Temperature considerations when working with GPMVs

With the following exception, all of the work presented thus far was carried about at 21-23 °C. Phase separation is highly temperature dependent and previous work found that the temperature at which half of GPMVs from HeLa cells phase separate is roughly 25 °C (24). The instrument used for most of this work is limited at low end of temperature to about 21°C. So, we could not easily probe the effect size of raft destabilizing compounds by further decreasing the temperature (which would increase phase separation). Our ability to heat samples was less restricted. In addition to the altered effects on raft stability in response to PD treatment, there was also a more modest increase in MAL ordered partitioning at 27 °C vs 23 °C (fig. S1D). This illustrates that temperature is a crucial variable to consider when interpreting the effects of these compounds on raft stability and ordered partitioning.

### Plate reader membrane fluidity assay

For fluidity measurements in GPMVs, GPMVs were made as described above without any staining prior to vesiculation. Once collected and settled, GPMVs were stained with Di-4-ANEPPDHQ (Invitrogen, cat # D36802). To optimize the concentration of each stain a pilot study was conducted with 0.1% DMSO and single compound and dye concentrations ranging from 0.5 µM to 10 µM. From this, 1 µM as determined to give the best effect sizes for Di-4. For experiments with all compounds GPMVs were treated with dye then deposited in wells of a 96-well plate with compounds (10 µM final for each compound) and incubated at room temperature for about 40 minutes. After incubation the plate was read on a SpectraMax iD3 plate reader (Molecular Devices). Data were acquired using SoftMax Pro 7 version 7.1.0 (Molecular Devices). Di-4 was excited at 470 nm and emission spectra were collected from 550 to 800 nm using 2 nm steps. The photomultiplier tube gain was set to automatic with an integration time of 140 ms.

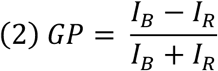

From the spectra, generalized polarization values were calculated with Equation 2. Where *I_B_* is a value in the blue end of the emission spectra and *I_R_* is a value at the red end. For Di-4 565 nm and 605 nm were selected for *I_B_* and *I_R_* respectively.

Similar experiments were conducted in ordered and disordered large unilamellar vesicles (LUVs) using previously described lipid mixtures (32). Briefly, powdered lipid stocks were purchased of cholesterol (Sigma, C3045), Egg Sphingomyelin (SM) (Avanti, 860061P), and 1-Palmitoyl-2-oleoyl-sn-glycero-3-phosphocholine (POPC) (Corden Pharma, LP-R4-031). 10 mM stocks were made in either chloroform (POPC and cholesterol) or ethanol (SM). Lipid films were dried down (L_o_: 25% POPC, 35% SM, 40% cholesterol, 70% POPC, 25% SM, 5% cholesterol) in glass culture tubes under a stream of nitrogen gas and further dried under vacuum for at least 2 hrs. Lipids films were resuspended in 50 mM HEPES, 150 mM NaCl, 0.2 mM EDTA pH 7.4 at 1 mM. These mixtures were then extruded (Avanti mini-extruder, 610000) through polycarbonate membranes with a 100 nm pore size (Whatman Nuclepore, 800309) on a heat block to warm the extruder block to ∼63 °C. LUVs were then diluted to 200 µM and labeled with 2 µM Di-4 and treated with compounds (or DMSO control) at a final compound concentration of 10 µM followed by a 45 minute incubation in a 96-well plate. Plates were then read on a plate reader and GP values calculated as described above.

### Image-based membrane fluidity assays

For imaging, GPMVs were prepared, treated with compounds, and labeled with Di-4 in the same manner as described in the previous section. They were seeded in an 8-well chamber slide with a coverslip on the bottom (Ibidi cat # 80806). Images were acquired on a Zeiss LSM 880 laser scanning confocal using a spectral detector and a 40X oil immersion objective. Images were collected at emission wavelengths from 410 nm to 689.5 nm at an interval of 8.9 nm.

Live-cell experiments were conducted by first seeding 10,000 cells per well in 8-well chamber slides. The following day, 1 hr prior to imaging, cells treated to deplete cholesterol were first rinsed with serum-free media then incubated with 10 mM methyl-ß-cyclodextrin (MBCD) in serum-free DMEM. After 30 minutes at 37 C, cells were removed from the incubation and treated with 10 µM compounds in serum-free CO_2_ independent media (Gibco L15 media, # 2108302) for 30 min. (MBCD treated well was also swapped from 10 mM MBCD in serum-free CO_2_ independent media). Prior to imaging, Di-4 was added to a final concentration of 2 µM. Imaging and Di-4 addition were staggered to ensure less than 30 min passed after addition of the dye and imaging. This is in line with previous observations that Di-4 begins to accumulate in endosomes after 30 min. Additionally, images were acquired at room temperature to slow internalization of the dye. Imaging was conducted on the LSM 880 as described in the previous section. Fluorescence intensities of individual cells were measured in ImageJ across all 32 wavelengths. 2 to 3 cells were measured using Fiji (68) from 5 fields of view per condition. GP values were calculated using equation 2 with intensities values for 561.5 nm and 605.8 nm used as the blue and red values respectively.

### Trypan blue cell viability

To ensure that effects seen in Di-4 live cell fluidity experiments and APC experiments were not due to inherent toxicity of the compounds, trypan blue experiments were conducted. HeLa cells were collected and treated with 10 µM compound or 10 mM MBCD as in the Di-4 assay. Live-dead staining was then conducted with trypan blue as previously described (69). Cells were mixed in a 50:50 ratio with trypan clue reagent, then immediately quantified on an automated Countess 3 cell counter (Fisher Scientific). Experiments were conducted on three separate days with measurements taken in duplicate.

### Automated patch clamp electrophysiology

HEK293 cells stably expressing full-length human TRPM8 were grown in DMEM media (Gibco 11960077) with 10% fetal bovine serum (Gibco 16000), 4 mM L-glutamine (Gibco 25030), 100 U/mL penicillin-streptomycin (Gibco 15140), 100 µM non-essential amino acid solution (Gibco 11140050), 4 mM GlutaMAX (Gibco 35050061), 200 µg/mL G418 (Sigma-Aldrich A1720), and 0.12% sodium bicarbonate (Gibco 25080094) at 37 °C and 8% CO_2_ in 100 mm dishes, as previously (70). After growing to 75% confluency (3-4 days), the cells were washed twice with 2 mL per dish of phosphate buffer saline solution (PBS), pH 7.4 (Gibco 10010031) followed by incubation of accutase (2 mL per dish, Gibco A1110501) for 5 minutes at 37 °C. Cells were then triturated and transferred to a conical tube and centrifuged at 200 ×g for 1.5 min to remove the accutase. The cells were resuspended with serum-free media (Gibco 11686029) and transferred into a T25 flask. The cells recovered for at least 30 mins at room temperature by gentle shaking (50 rpm). Following recovery, the cells were centrifuged (200 x g for 1.5 minutes) and resuspended in extracellular buffer (10 mM HEPES, 145 mM NaCl, 4 mM KCl, 1 mM MgCl_2_, 2 mM CaCl_2_, 10 mM glucose, pH 7.4) to a cell density of 3-7 × 10^6^ cells/mL. The osmolality of the extracellular buffer was adjusted using a Vapro 5600 vapor pressure osmometer (Wescor) with sucrose to 315-330 mOSm and pH using NaOH.

Data was collected using IonFluxMercury HT (Cell Microsystems) automated patch clamp electrophysiology instrument with Ionflux HT v5.0 software using ensemble IonFlux Plate HT (Cell Microsystems 910-0055). The ensemble microfluidic plates enable 32 parallel experiments with aggregate currents from 20 cells per experiment. Intracellular solution was composed of 10 mM HEPES, 120 mM KCl, 1.75 mM MgCl_2_, 5.374 mM CaCl_2_, 10 mM EGTA, 4 mM NaATP, pH 7.2. The intracellular solution osmolality was adjusted with sucrose to 305-315 mOsm and the pH was adjusted using KOH. Compounds were dissolved into DMSO before adding to extracellular solution, where the DMSO concentration was kept consistent at 0.03% v/v across all compounds and controls. Prior to experiments the plates were washed as suggested by the manufacturer. The protocol for the experiment is divided into four steps: prime (priming microfluidics with solutions), trap (trap cells and obtain membrane seals), break (access to intracellular by breaking membrane), and data acquisition. Each step also has multiple channels: main channel (positive pressure allows solutions to flow towards the cells and waste), trap channel (negative pressure to provide a vacuum to keep the cells in the traps and break the cell membrane to access intracellular), and compound channel (positive pressure to flow compounds to main channel). During the prime step: (1) the main channel was applied 1 psi for t = 0-25 s and 0.4 psi for t = 25-60 s (2) the trap and compound channels were applied at 5 psi for t = 0-20 s and then 1.5 psi for t = 20-55 s followed by only the traps at 2 psi for t = 55-60 s. For the voltage during the prime step, a pulse was applied every 150 ms, where the 0 mV holding potential was applied during t = 0-50 ms, 20 mV was applied during the t = 50-100 ms, and 0 mV during the t = 100-150 ms. During the trap step: (1) the main channel is applied 0.1 psi for t = 0-5 s before applying 0.5 s pulses of 0.2 psi every 5 s during t = 5-135 s (2) the trap channel is applied 6 inHg for t = 0-135 s. For the voltage during the trap step, a pulse was applied every 70 ms, where the -80 mV holding potential was applied between t = 0-20 ms, -100 mV for t = 20-50 ms, -80 mV for t = 50-70 ms. During the break step: (1) the main channel is applied 0.1 psi for t = 0-100 s (2) the trap channel was applied 6 inHg between t = 0-10 s, vacuum ramp from 10 to 14 inHg from t = 10-40 s, and 6 inHg for t = 40-100 s. For the voltage during the break step, a pulse was applied every 150 ms, where -80 mV holding potential was applied between t = 0-50 ms, -100 mV for t = 50-100 ms, and -80 mV for t = 100-150 ms. During the data acquisition: (1) the main channel is applied at 0.15 psi for t = 0-1350 s and 0 psi for 1350-2450 s, (2) the traps channel is applied 5 inHg for t = 0-3 s, 3 inHg for t = 3-1350 s, and 0 inHG for t = 1350-2450 s. For the voltage during the data acquisition, a pulse was applied every 625 ms, where the -60 mV holding potential was applied between t = 0-100 ms, -70 mV for t = 100-200 ms, -60 mV for t = 200-300 ms, a voltage ramp from - 120 mV to 160 mV for t = 300-525 ms and -60 mV for t = 525-625 ms. The cells/plates were equilibrated to 30 °C for 5 minutes. Prior to application of compounds, menthol was perfused for 75 s four times to measure initial current responses. Compounds were then applied by continuous perfusion for 15 minutes followed by measurement of two 75 s applications of menthol in the presence of compound.

Ionflux Data Analyzer v5.0 was used to analyze the data. Leak subtraction was performed on the data based on the -60 mV initial holding potential and -70 mV voltage steps from the data acquisition. Each point of the current trace is from the difference of the current at 120 mV and the holding potential at -60 mV. The data was averaged from 7 points after 25 s of perfusion of menthol without or with the compound. The last two menthol stimulated currents before compound application were averaged and compared to the two menthol-stimulated currents after compound application to determine a ratio of menthol response.

### TRPM8 cell surface measurements

TRPM8 stable cells were cultured as described above. Cells were collected by dissociation with 0.5 mM EDTA in PBS and resuspended in media. Cells were then incubated in 100 µl of media with 10 µM compound or 10 mM MBCD for 15 minutes as in the electrophysiology experiments. Cells were then fixed with 100 µl Buffer A from a Fix & Perm kit for flow cytometry (Invitrogen, Cat. No. GAS004). Cells were then rinsed 3 times in flow cytometry buffer (PBS + 5% FBS + 0.1% NaN_3_). Cells were then labeled with either of two TRPM8 primary antibodies targeted to an extracellular epitope (Alomone, Cat. No. ACC-049 Abcepta, Cat. No. AP8181D). The Alomone antibody was used at a dilution of 1:100 while the Abcepta antibody was used at a dilution of 1:50 for 1 hr in the flow cytometry buffer. Cells were rinsed 3 times again then labeled with an anti-Rabbit-AlexaFlour488 secondary at a 1:1000 dilution (Cell Signaling, Cat. No. 4412) for 45 min. Cells were rinsed 3× again and resuspended in a final volume of 300 µl. Single cell fluorescence intensities were measured on a BD Fortessa 5-laser analytical cytometer. Geometric means of the resulting intensity distributions were calculated in FlowJo (version 10). Statistical comparisons were made in GraphPad Prism (version 10).

### Immunoblotting to detect EGFR activation

For EGFR activation studies, adherent HeLa cells at 70 % confluency in a 60 mm dish were starved overnight (∼18 hr) with starvation media - serum free DMEM/F12 (Gibco) media supplemented with only Pen-Step. Starved cells were exposed to small molecules of interest by replacing the overnight starvation media with 4 mL of pre-warmed starvation media containing 10 µM of compounds of interest and 0.1 % DMSO (Cell Signaling Technologies) for noted times at 37 °C. After incubation, EGFR was activated with the addition of 1 mL of pre-warmed starvation media containing 500 ng/mL EGF (R&D Systems) for 1 minute. Due to the speed of EGFR activation kinetics in HeLa cells, treated plates were then flash frozen in liquid nitrogen after media removal. Frozen plates were then placed on ice and lysed with scraping in ice-cold RIPA lysis buffer supplemented with PhosStop phosphatase inhibitor and Complete protease inhibitor (Roche). Lysates were clarified by centrifugation and subjected to immunoblotting using NuPage Novex 4 % - 12 % Bis-Tris Protein Gels (ThermoFisher Scientific). After electrophoresis, intact gels were transferred to Immoblion-P PVDF (Millipore) membranes and cut into three horizontal strips guided by Precision Plus molecular weight ladder (Bio-Rad) and incubated overnight in blocking buffer - 20 mM Tris, 150 mM NaCl, 0.1 % Tween-20 (Bio-Rad) pH 7.6 (TBST) with 3 % w/v Bovine Serum Albumin Fraction V (Fisher Bioreagents). Primary rabbit antibodies against EGFR pY1173 (53A5, 3972S), phospho-p44/42 MAPK – also known as ERK 1/2 (9101S), and GRB2 (3972S) were purchased from Cell Signaling Technology and used at a dilution of 1:1000 in TBST for 1 hr at RT with gentle agitation. Goat anti-rabbit IgG (H+L) conjugated to horse radish peroxidase (ThermoFisher Scientific, 31460) with glycerol was used at a dilution of 1:5000 for 1 hr in blocking buffer with gentle agitation. Blots were detected using SuperSignal West Pico Chemiluminescent Substrate (ThermoFisher Scientific) on a LI-COR 2800 using the chemiluminescent and 700 nm channels to detect antibody and molecular weight bands signals, respectively. Chemiluminescent signal was checked for saturation and bands of interest were integrated with Image Studio (LI-COR, version 3.1).

### Statistical Comparisons

All Statistical comparisons were conducted in GraphPad Prism (version 10.3.1). Where multiple comparisons are meant to be interpreted together, ANOVA were conducted followed by the Dunett’s test for pairwise comparisons or the non-parametric Mann-Whitney test. Where stand-alone comparisons of two means that are not meant to be interpreted with other comparisons are made, student’s t-tests are used. Only P-values of less than 0.05 are considered significant.

## Acknowledgments

The MAL-GFP construct was a gift from the Levental lab (University of Virginia). Some experiments were performed in the Vanderbilt High-Throughput Screening (HTS) Core Facility with assistance provided by Corbin Whitwell. The FDA approved library was provided by the Vanderbilt CTSA and distributed by the Vanderbilt High-Throughput Screening Core Facility as was the Vanderbilt Discovery Collection. The HTS Core receives support from the Vanderbilt Institute of Chemical Biology and the Vanderbilt Ingram Cancer Center.

Membrane fluidity experiments were performed in part through the use of the Vanderbilt Cell Imaging Shared Resource.

## Author contributions

Conceptualization: KMS, AKK, WDHV, CRS

Methodology: KMS, HH, JAB, AKK, WDVH, CRS

Investigation: KMS, HH, DDL, JMH, NS, AJF, TPH

Supervision: JAB, AKK, WDVH, CRS

Writing—original draft: KMS, CRS

Writing—review & editing: KMS, HH, DDL, JAB, AKK, WDVH, CRS

## Data and materials availability

All data are available in the main text or the supplementary materials.

