## Supplemental Materials for "Pharmacological Tools to Modulate Ordered Membrane Domains and Order-Dependent Protein Function"

**This File Includes:**

**Supplementary Figures S1-S6**

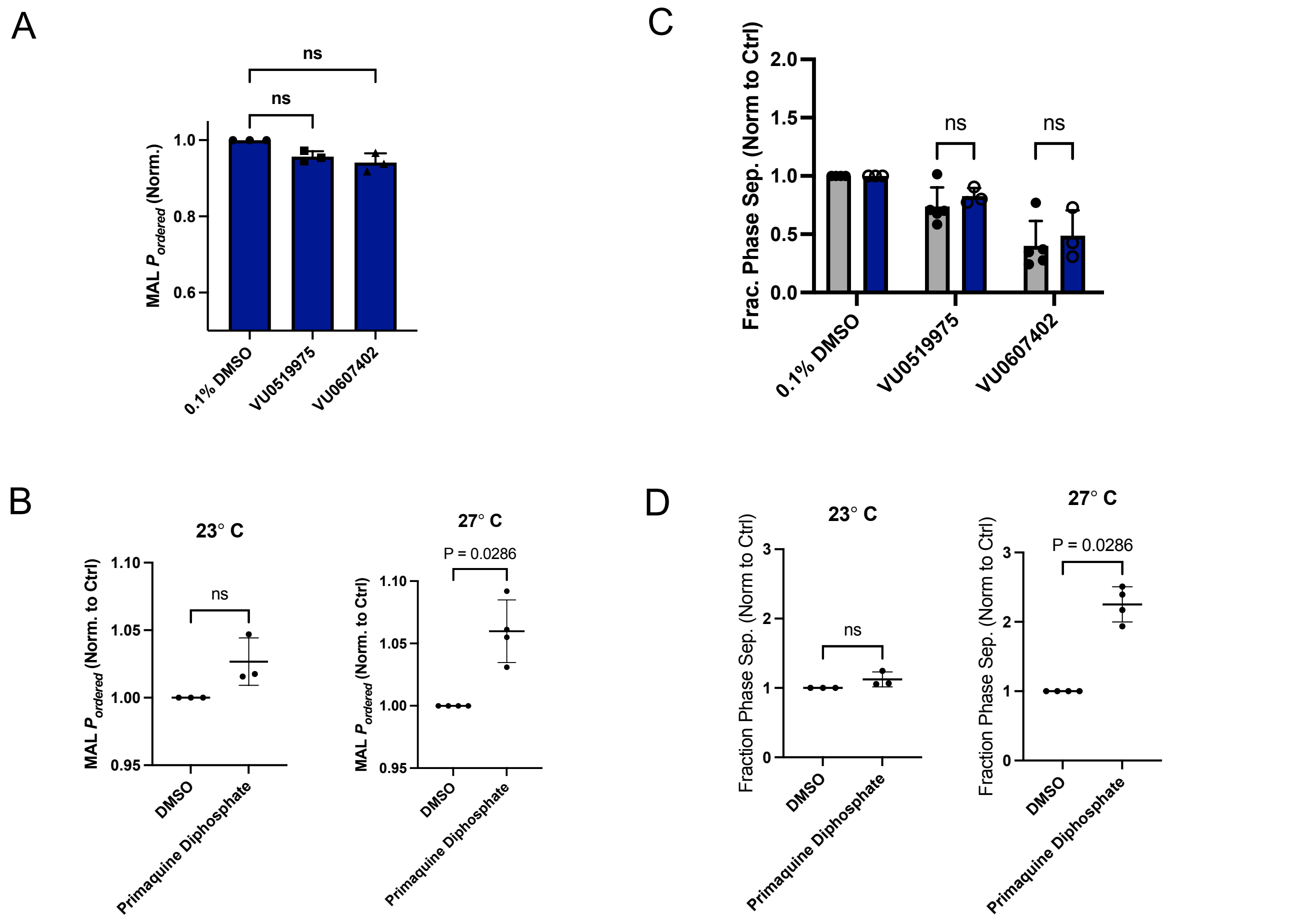


**Figure S1. Compounds effects on $P_{ordered}$ and raft stability in GPMVs expressing MAL. A)** Effects of 10 µM protein-independent hits on ordered partitioning of MAL. Bars are means ± SD (*n* = 3). **B)** Effects of 10 µM primaquine diphosphate on ordered partitioning of MAL at two temperatures. Bars are means ± SD (*n* = 3). **C)** Effects of 10 µM hits on phase separation with (navy bars) and without (gray bars) expression of MAL (untransfected cells). Bars are means ± SD (*n* = 3-5). **D)** Effects of 10 µM primaquine diphosphate on phase separation at two temperatures. Bars are means ± SD (*n* = 3-4). P-values are from Mann-Whitney tests.

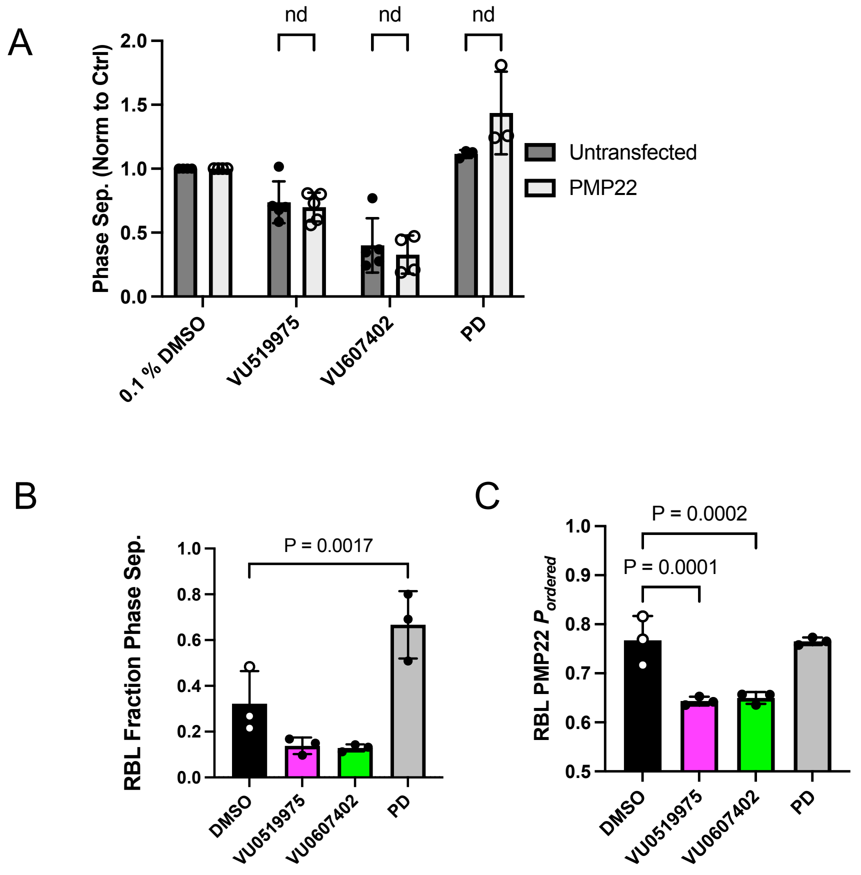


**Figure S2. Effects of compounds on both phase separation and PMP22 ordered partitioning are similar in RBL cells as compared to HeLa and effects on phase separation are independent of PMP22 expression. A)** Effects of 10 μM hits on phase separation with (light gray) and without (dark dray) expression of PMP22. Bars are means ± SD (*n* = 3-5). **B)** At 5 μM, compounds that are raft destabilizing in GPMVs from HeLa cells also decrease the fraction of phase separated GPMVs from RBL cells, though the change is not statistically significant likely due to the low starting levels of phase separation in RBL GPMVs. Primaquine diphosphate significantly increases the fraction of phase separated GPMVs. n = 3, bars are means ± SD. Statistical comparisons are from ANOVA followed by Dunnett's tests. **C)** 5 μM ordered domain destabilizing compounds decrease PMP22 ordered partitioning in GPMVs derived from RBL cells.

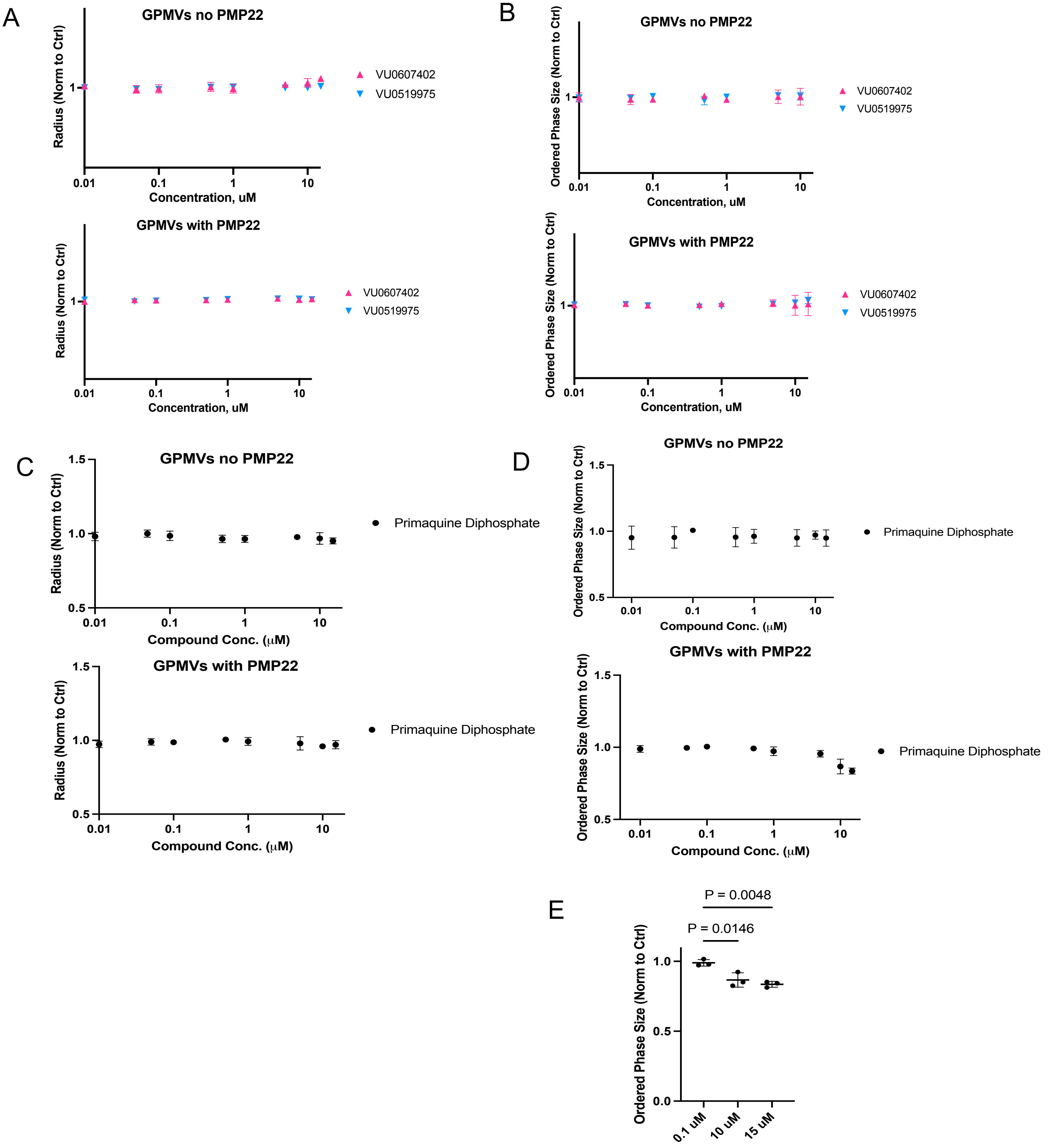


**Figure S3. Effects of compounds on HeLa derived GPMV size and size of ordered domains with and without PMP22 expression. A)** Average radii and relative size of ordered domains of GPMVs from untransfected cells treated with hit compounds across a range of doses normalized to DMSO control. **B)** Average radii and relative size of ordered domains of GPMVs from PMP22 expressing cells treated with hit compounds across a range of doses normalized to DMSO control. **C)** Average radii of GPMVs -/+ PMP22 treated with primaquine diphosphate. **D)** Average ordered domain size in GPMVs -/+ PMP22 treated with primaquine diphosphate. **E)** Comparison of change in domain size seen in GPMVs with PMP22 treated with PD. Points and bars are means ± SD. P-values are from ANOVA followed by Dunnetts tests.

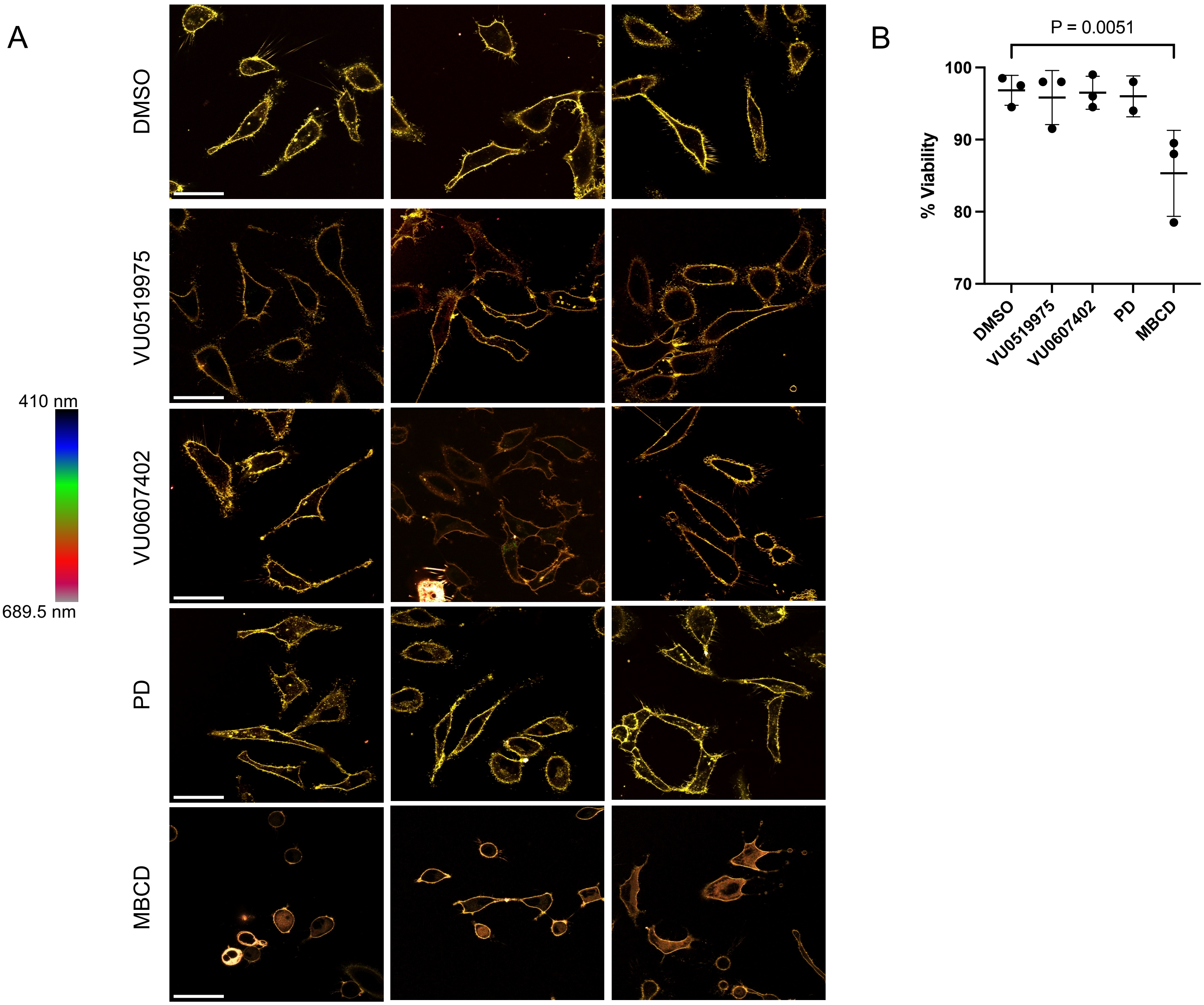


**Figure S4. Representative results of live cell membrane fluidity experiments and corresponding viability experiments. A)** Three representative spectral images from each compound treatment used in Fig 5 D and E. Cells labeled with Di-4. Scale bars are 50 µm, scale is identical for each panel. **B)** Trypan blue viability experiments conducted with HeLa cells treated with compounds as they were in the Di-4 microscopy experiments. N =3, bars are means ± SD, P-value is from ANOVA followed by Dunnett's test.

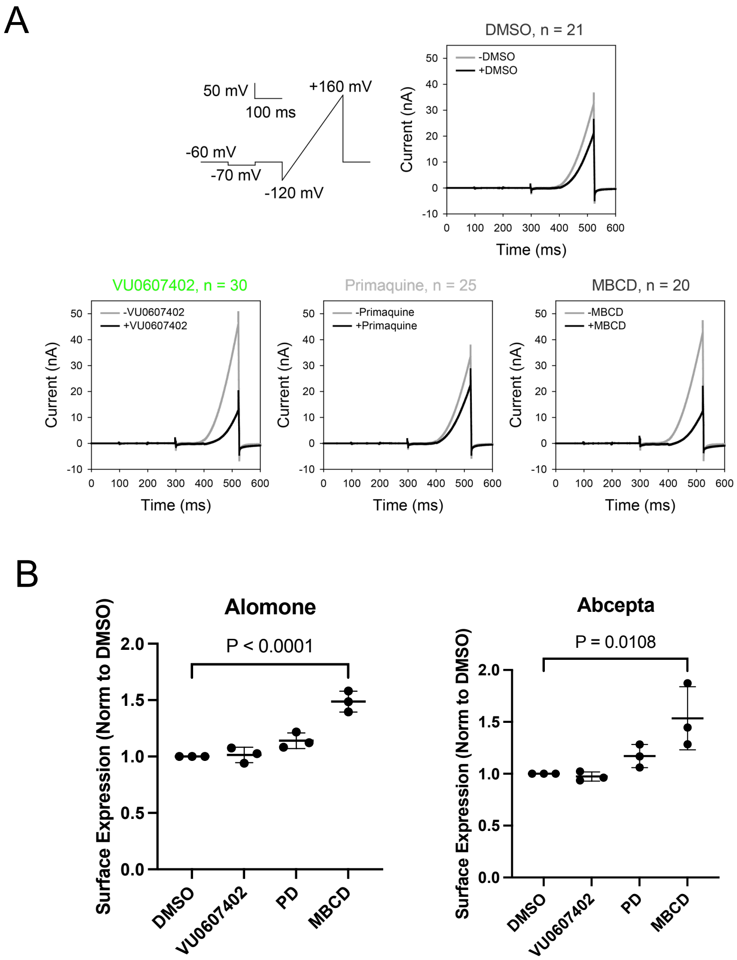


**Figure S5. Effects of compounds on TRPM8 activity and cell surface expression. A)** The average current pulse of TRPM8 before and after exposure to the compounds. The top-left shows a schematic of the pulse program. Graphs are current response against time from a single pulse program at 922.2 seconds and 1971.99 seconds from panel A the 100 µM menthol response without (-) and with (+) 0.03% DMSO (control), 10 µM VU0607402, 10 µM PD, or 10 mM MBCD. Each n refers to the single sum of 20 cells from the plate **B)** Quantification of plasma membrane TRPM8 in stable cells by flow cytometry using two different extracellularly directed TRPM8 antibodies following 15 min treatment with each compound as in the APC experiments. n =3 bars are means ± SD. P-values are from ANOVA followed by Dunnett's tests.

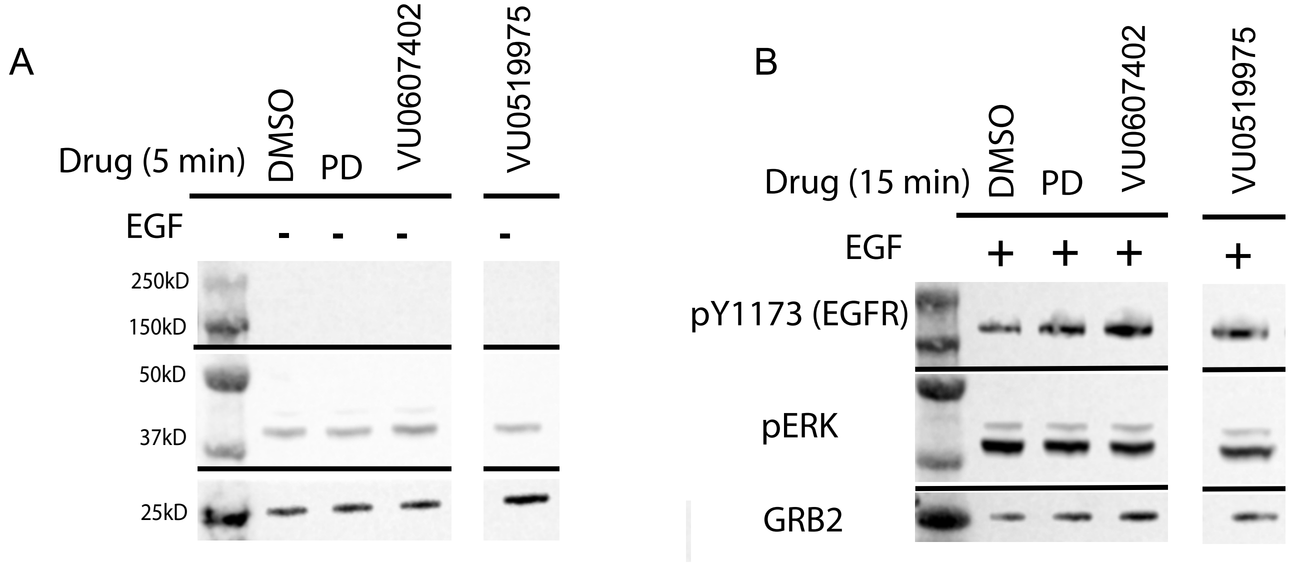


**Figure S6. Representative Western blot of phospho- EGFR and ERK.** A) Representative blot of HeLa cell lysates from cells incubated with compounds for 5 minutes and not stimulated with EGF B) Representative blot of HeLa cell lysates incubated with compounds for 15 min then stimulated with EGF for 1 min.
